# Reduced cortico-muscular beta coupling in Parkinson’s disease predicts motor impairment

**DOI:** 10.1101/2021.05.15.444289

**Authors:** Nahid Zokaei, Andrew J Quinn, Michele T Hu, Masud Husain, Freek van Ede, Anna Christina Nobre

**Affiliations:** Oxford Centre for Human Brain Activity, Wellcome Centre for Integrative Neuroimaging, Department of Psychiatry, University of Oxford, Oxford, OX3 7JX; Department of Experimental Psychology, University of Oxford, Oxford, OX2 6GG; Department of Psychiatry, University of Oxford, Oxford, UK, OX3 7JX; Department of Neurology, Nuffield Department of Clinical Neurosciences, University of Oxford, Oxford, UK, OX3 9DU

## Abstract

Long-range communication through the motor system is thought to be facilitated by phase coupling between neural activity in the 15-30 Hz beta range. During periods of sustained muscle contraction (grip), such coupling is manifest between motor cortex and the contralateral forearm muscles – measured as the cortico-muscular coherence (CMC). We examined alterations in CMC in individuals with Parkinson’s disease (PD), while equating grip strength between individuals with PD (off their medication) and healthy control participants. We show a marked reduction in beta CMC in the PD group, even though the grip strength was comparable between the two groups. Moreover, the reduced CMC was related to motor symptoms, so that individuals with lower CMC also displayed worse motor symptoms. These findings highlight the CMC as a simple, effective, and clinically relevant neural marker of PD pathology, with the potential to aid monitoring of disease progression and the efficacy of novel treatments for PD.

## Introduction

Parkinson’s disease (PD) is a neurodegenerative disorder characterized by a loss of dopaminergic neurons that alters neural activity in the basal ganglia, thalamus, and the cortex (Lang & Lozano, 1998). Excessive activity in the beta band (13-30 Hz) in the basal ganglia is a hallmark of PD (see Hammond et al., 2007 for a review). Local field-potential (LFP) recordings from subthalamic nucleus of individuals with PD also show excessive beta activity, which is reduced following dopaminergic medication (Brown et al., 2001; Kühn et al., 2004; Levy et al., 2002; Priori et al., 2004) or deep brain stimulation (DBS) treatment (Eusebio et al., 2012; Giannicola et al., 2010; Kühn et al., 2008).

Contrary to the excessive beta activity in the basal ganglia, cortical beta in individuals with PD may be attenuated instead (Bosboom et al., 2006; Heideman et al., 2020; Stoffers et al., 2007; Vardy et al., 2011). Moreover, prior studies have suggested that neural coupling between the cortex and the muscle – measured as the cortico-muscular coherence (CMC) – may also be reduced in PD (e.g. Airaksinen et al., 2015; Hirschmann et al., 2011; Park et al., 2009; Pollok et al., 2012; Salenius et al., 2002). The CMC quantifies the temporal relationship (phase coupling) between neural activity at a given frequency in the motor cortex and in the contralateral muscles, and is particularly pronounced during periods of steady muscle contraction (Baker, 2007; Tatsuya Mima et al., 2002; Salenius et al., 1997; Schoffelen et al., 2005). It is also impacted by other disorders involving motor impairments, such as stroke (Farmer et al., 1993; Nielsen et al., 2008; Rossiter et al., 2012) or amyotrophic lateral sclerosis (ALS) (Fisher et al., 2012; Proudfoot et al., 2018), and may be used to track disease progression and therapeutic success (Airaksinen et al., 2015; von Carlowitz-Ghori et al., 2014; McKeown et al., 2006; Salenius et al., 2002).

However, previous studies on changes in CMC in PD have limitations. Firstly, the required motor output during the period at which CMC was calculated, has typically not been carefully equated between PD and healthy-control groups, making it difficult to disambiguate if differences in CMC were specific markers of disease or simply related to secondary differences in motor output. In addition, sample sizes were small in most prior studies. In many cases, measurements may also have been contaminated by concurrent DBS effects.

Here, we aimed to overcome these shortcomings by revisiting CMC in individuals with PD in a task using carefully controlled and well equated grip strength between a group of 17 individuals with PD who were off medication and 17 matched healthy controls. We show that CMC is markedly reduced in individuals with PD even when grip force was equated. Importantly, CMC strength also predicted motor impairments.

## Methods

### Participants

The study was approved by the Oxford Research Ethics Committee as part of the National Research Ethics Service, and participants gave written informed consent to task procedures prior to participation.

Seventeen individuals with PD and seventeen age- and education-matched healthy control participants took part in this study (**Table 1** for demographics). Participants in the PD group were recruited from neurology clinics in Oxfordshire (UK). Exclusion criteria were: being an active participant in an ongoing clinical drug trial, not tolerating coming off medication, taking psychotropic hypertensive or vasoactive medication or long acting dopamine agonists, or a history of neurological or psychiatric disease other than PD. Healthy control participants were recruited via the Friends of OxDare registry (https://www.oxdare.ox.ac.uk/become-a-friend). All participants had normal or corrected-to-normal vision.

Participants with PD were asked to withdraw from their dopaminergic medication at 7PM the night before the experiment. The Addenbrooke’s Cognitive Examination (ACE |||) test was administered as a general cognitive screening test to the PD and healthy-control groups (**Table 1**). Even though individuals with PD scored significantly lower than healthy individuals, no participant had generalized cognitive impairment (as set by a cut-off point of 85). Unified Parkinson’s Disease Rating Scale (UPDRS) was administered prior to the scanning to all participants with PD.

**Table 1.** Demographics, Addenbrookes Cognitive Examination-III (ACE-III) and UPDRS section III scores of PD and healthy-control participants.

| | PD participants | | Control participants | | $\Delta$ |
| --- | --- | --- | --- | --- | --- |
|  | Mean (SE) | Range | Mean (SE) | Range | *P* |
| Age (years) | 66.5 (1.2) | 52 - 78 | 69.7 (2.01) | 54 - 80 | n.s. |
| Education (years) | 17.2 (1.5) | 10 - 24 | 16.7 (0.9) | 10 - 20 | n.s. |
| ACE – III | 93.9 (4.03) | 88-100 | 96 (3.3) | 90-100 | 0.044 |
| UPDRS section III | 29.4 (3.23) | 11-44 | n/a | n/a | n/a |
| Daily Levodopa Equivalent dose | 308 (135.7) | 150-540 | n/a | n/a | n/a |

### MEG acquisition

Magnetoencephalography (MEG) data were acquired using an Elekta Neuromag system with 306 channels, at a sampling rate of 1000 Hz. Participants were seated comfortably in the MEG scanner and performed the task after being familiarized with the task procedure. Head position was tracked continuously using 4 fixed coil positions relative to the nasion and pre-auricular fiducial landmarks. The Electrocardiogram (ECG) was monitored by placing electrodes at both wrists. Saccades and blinks were monitored using recordings from electrodes around one of the eyes to derive the horizontal and vertical electro-oculogram, respectively. Muscle contraction was measured using bipolar electromyography (EMG) recordings at both forearms using electrodes placed approximately 4 cm apart over the flexor digitorum superficialis of each arm, with reference electrodes on the lateral epicondyle (as in Proudfoot et al., 2018).

### Grip task and procedure

**Figure 1a** contains a schematic of the grip task. In each trial, participants were presented with two bars on either side of the fixation cross corresponding to each hand (500 msec). The force to be exerted (target force) was then indicated by the height of two red lines on each bar (matched between the two hands). Across trials, participants had to produce and maintain a light grip of approximately 12 Newtons or a strong grip of approximately 17 Newtons. Using the same gripper devices, previous studies have reported no difference in maximum voluntary contraction (MVC) in individuals with PD and healthy individuals (Chong et al., 2015). Participants had to exert and maintain the grip for 3 seconds, after which time the red lines dropped to the bottom of both bars. Grip strength was recorded using MEG-compatible fibre-optic auxotonic-force devices or “grippers” (Current Designs, USA). Participants received direct visual feedback on the force they exerted on each gripper (**Figure 1a**, blue bars). Participants could relax for 2000 ms between successive grip trials. All participants performed 120 trials of the task (60 per force condition) across 12 blocks of 10 trials each.

Some participants across both PD and HC groups had difficulty reaching and/or maintaining the required stronger grip force in many trials. We therefore restricted our analyses to the light-grip trials, which all participants could comfortably manage. Both groups successfully completed the task in time, and none of the participants reported being fatigued by task procedure.

**Figure 1.**
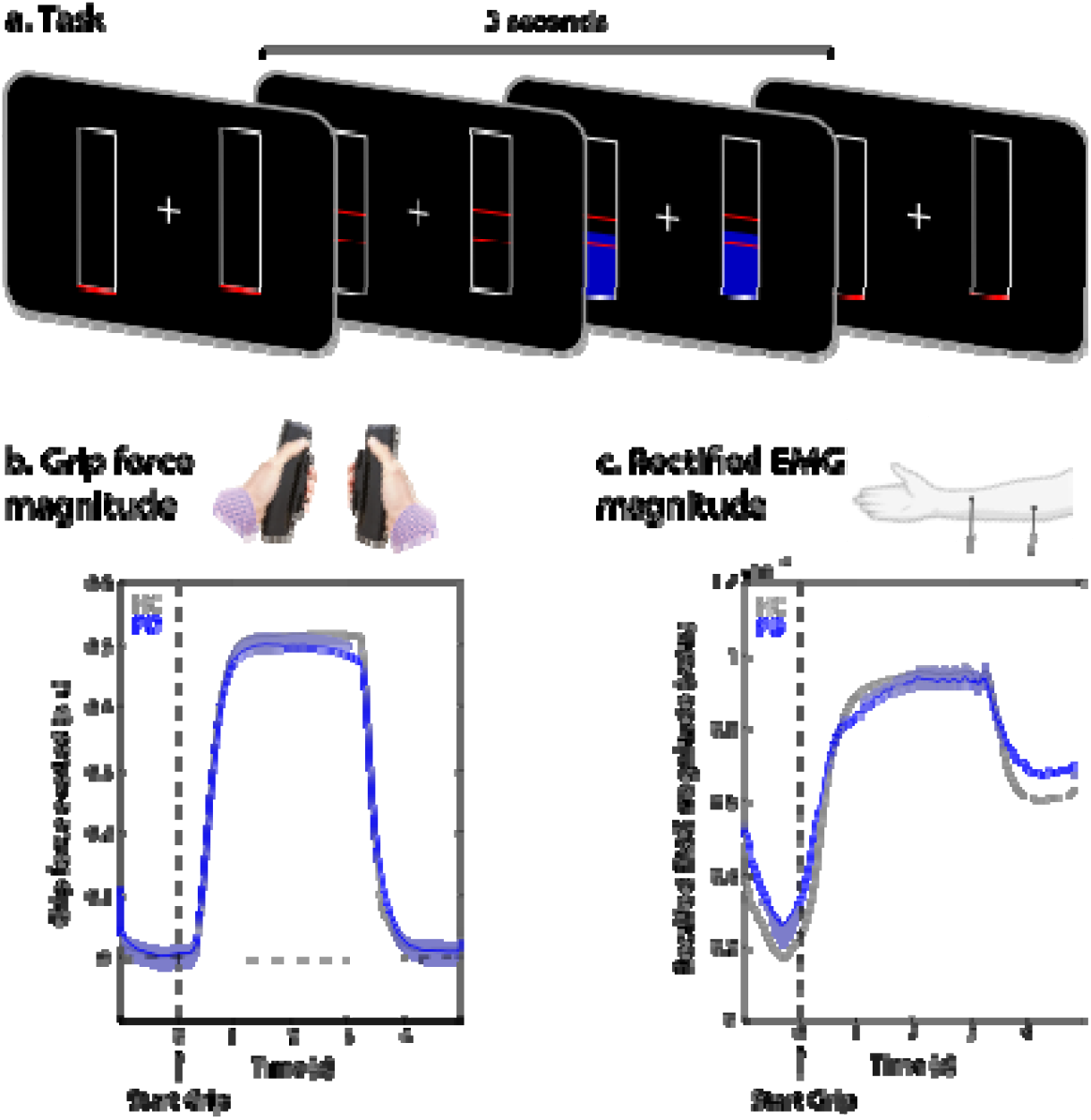
a) Schematic of the grip task in which participants had to exert bilateral auxotonic grip (isometric contraction) for a duration of 3 seconds. b-c) The amount of force exerted was similar between healthy-control (grey) and PD (blue) participants, as measured in gripper output (b) and rectified EMG (c). The shaded grey areas indicate the period of “stable grip” used in the analysis of CMC.

Trials in which participants clearly did not reach or maintain the required grip level (in which the force exerted reached zero during the grip duration, after baseline correction) were excluded from analysis. Both PD and HC participants could perform the task successfully, with an average of 56 (out of 60) trials completed by HC participants (SD: 3.12) and 56.7 trials by PD participants (SD: 4.2). There was no significant difference between the number of usable trials completed by HC and PD participants (*t*(32)= 0.61, *p*= 0.5).

### Preprocessing

Spatial signal space separation and movement compensation were applied using Neuromag’s Maxfilter software to separate signals arising from inside vs. outside of the MEG helmet (minimizing extracranial noise) and compensate for the effects of head movements using continuous head-position measurements. Data were then converted to fieldtrip format, checked manually to ensure no problems arose during the Maxfilter pre-processing stage, and downsampled to 250 Hz. Low frequency drift was removed by using a 0.1-Hz high-pass filter. The down-sampled data were epoched; and Independent Component Analysis (ICA) was used to remove artifacts associated with blinks, saccades, and heartbeat. Any remaining artifacts were rejected following visual inspection of the data. To investigate muscle contraction during the grip period, we additionally constructed a channel with high-pass filtered (40-Hz cut-off) and baseline-corrected EMG traces (averaged between the left and right arm).

### Corticomuscular coherence estimation

The CMC is a measure of phase-coupling (normalized to range from 0 to 1) between the cortical motor MEG signal and the corresponding contralateral muscular EMG signal that is particularly prevalent during continuous, isometric, contraction (Baker, 2007; T. Mima & Hallett, 1999; Proudfoot et al., 2018; Salenius et al., 1997; Schoffelen et al., 2005). We used the rectified raw EMG traces when calculating CMC. CMC was calculated using the FieldTrip toolbox (Oostenveld et al., 2011), using the data from the steady contraction window between 1000 and 3000 ms after the onset of the grip instruction. CMC was calculated between 1 to 40 Hz, in 0.25 Hz steps. A multi-taper method (Percival & Walden, 1993) was applied to achieve ±5 Hz spectral smoothing. We focused our analysis of CMC magnitude on MEG signals from the 12 combined planar-gradiometer channels (6 per hemisphere) which a previous, independent, study from our lab showed to be particularly sensitive to motor cortical activity (4). We used the same set of left and right channels for both PD and HC groups. To avoid issues with multiple comparisons, we contrasted the CMC spectra between groups using a cluster-based permutation analyses (Maris & Oostenveld, 2007), with 5000 permutations.

## Results

We first ascertained that both the HC and the PD group had similar performance on the grip task (**Figure 1a**). As shown in **Figure 1b**, participants demonstrated a clear modulation of grip strength during the instructed grip period. Importantly, both groups achieved similar levels of grip force. We found no signification differences in mean grip force (**Figure 1b**) or in muscle contraction as measured through rectified EMG signals (**Figure 1c**) during the time of the grip (no significant clusters).

We next turned to our primary outcome measure of interest: cortico-muscular phase coupling, CMC. In line with prior studies of CMC (Baker, 2007; T. Mima & Hallett, 1999; Proudfoot et al., 2018; Salenius et al., 1997; Schoffelen et al., 2005), we observed particularly pronounced coupling between sensors over motor cortex and the contralateral muscle in the beta band. Frequency profiles were similar between groups. Taking the coupling with the ipsilateral muscle as our neutral “baseline”, we found significant contralateral (vs. ipsilateral) coupling in both PD (13-33 Hz, cluster *p*<0.0001; **Figure 2a**, blue line) and HC (7-27 Hz, 29-39 Hz, cluster *p*<0.0001; **Figure 2a**, grey line) participants. Critically, however, despite similar grip strength (**Figure 1b**), individuals with PD demonstrated a marked reduction in beta coupling between the cortex and contralateral muscle, as confirmed by a significant group difference (cluster-based permutation-significant cluster: 11-25 Hz, cluster *p*= 0.008; **Figure 2a**, black line). Moreover, there was no difference in CMC in the affected versus the unaffected side in individuals with PD (**Figure S1**, in fifteen individuals with unilateral symptom profile).

The reduction of CMC between cortex and contralateral muscle was also evident when considering its topographical distribution (**Figure 2b**). In both groups, coherence was localized to the same contralateral motor channels, but was considerably weaker in PD compared to HC participants.

**Figure 2.**
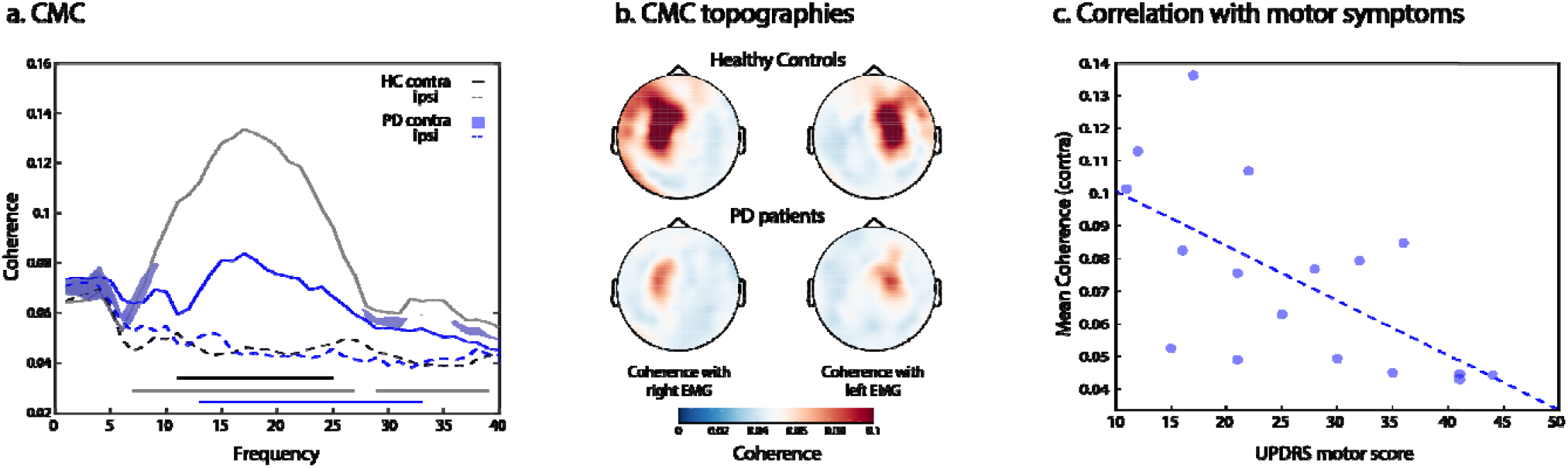
a) Cortico-muscular coherence (CMC) between the forearm EMG and contralateral (solid lines) and ipsilateral (dashed lines) motor cortex in the 1-3 s stable-grip period (shaded areas in Fig. 1b-c). Contralateral beta CMC was significantly reduced in individuals with Parkinson’s disease (blue line) compared to healthy control participants (grey line). Contralateral CMC was also significantly different between groups (black line). b) Topographical distributions of CMC in both participant groups, separately for coupling with the left and right EMG. c) Mean contralateral beta CMC, averaged over the significant cluster in a (11-25 Hz) was significantly correlated with motor symptoms in the PD group, as measured by the UPDRS.

We tested whether the reduction in contralateral beta coupling in PD compared to HC participants might be a relevant clinical marker. Within the PD group, the magnitude of CMC was negatively correlated to motor symptoms as measured by the UPDRS section III (motor symptoms) (**Figure 2c**). Lower (“more reduced”) coupling was associated with more severe symptoms (r=-0.63, *p*= 0.007).

In addition to robust group differences in cortico-muscular coupling, there were also significant group differences in cortical power (**Figure S2a;** with one cluster between 6 and 8 Hz, cluster *p*= 0.05, and another between 15.5 and 37 Hz, cluster *p*= 0.0019), as well as increased temporal *variability* in the rectified EMG (likely a reflection of tremor **Figure S2b**). Variability was measured by calculating variance in the rectified EMG, across 200 ms window that we advanced over the data in steps of 50ms. Critically however, these variables did not predict CMC or UPDRS section III scores (**Figure S3**). Moreover, when taking these variables into account, by partialling out mean power and EMG variability, the correlation between CMC and motor symptoms of UPDRS section III scores remained significant (*r*= −0.61, *p*= 0.015). Finally, there was no significant relationship between CMC and years of diagnosis (*r*= 0.15, *p*= 0.59) or daily levodopa-equivalent dose (*r*= −0.12, *p*= 0.68).

## Discussion

The results of this study demonstrated a marked reduction in beta CMC during a period of controlled sustained grip in the PD group, despite comparable grip magnitude to that of a matched healthy-control group. Moreover, the reduced CMC was related to motor symptoms in the PD group as measured by the UPDRS section III, so that individuals with lower CMC in the beta range also displayed worse motor symptoms.

Remarkably, the reduced CMC in individuals with PD did not impact their ability to perform the grip task in any obvious way. Even though the precise functional role of CMC is not fully understood (van Wijk et al., 2012), it has been shown to relate to both the quality and precision of motor performance as well as to skill learning in healthy participants (Kilner et al., 2000; Kristeva et al., 2007; Perez et al., 2006). Yet, the current work presents a curious situation in which a clear reduction in CMC occurred despite largely preserved grip performance. Importantly, the low magnitude of the required grip in our task was sufficient to reveal robust group differences in CMC while minimizing changes to overall motor performance. This advantageous context enabled us to re-evaluate changes in CMC in PD vs. HC, while minimizing the contribution of any group differences attributable to secondary differences in motor behaviour.

Deciphering the exact neurological mechanisms that underlie the observed group differences – and their link to disease severity – remains an important target for future studies. Previous studies have suggested that reduced CMC in PD may arise from deficits in programming of movement that is reflected in the loss of synchronized oscillatory activity in muscle discharge (Brown & Marsden, 1998; Marsden et al., 2001). More specifically, it has been hypothesized that CMC loads on the same pathways that are relevant and affected in bradykinesia (Brown, 2006). In addition, proprioceptive processing has been proposed to contribute to CMC(Baker, 2007) (Baker, 2007) and may also contribute to the observed differences.

Motor output (such as tremor present in participants with PD) also provides continuous motor/sensory input and could thereby reduce beta oscillations and as consequence result in a reduction in CMC. This intriguing possibility warrants further testing. However, attenuation of CMC by input associated with tremor was unlikely to account for our findings. In individuals with lateralized tremor in the PD group, the effects were equivalent when analyzing CMC in relation to the hand affected and unaffected by tremor (supplementary material S1). Moreover, when we regressed out participant-specific EMG variability – as a proxy for tremor – during the gripping period, we found that the relationship between clinical symptoms and CMC persisted.

Our findings build on related previous studies (e.g. Airaksinen et al., 2015; Hirschmann et al., 2011; Park et al., 2009; Pollok et al., 2012; Salenius et al., 2002), while also controlling for important prior limitations. Firstly, we measured CMC during a period of steady muscle contraction that was comparable between PD and HC participants. In most prior studies, participants were usually asked simply to extend their wrist or to contract their forearm, with no objective measure on the strength of muscle contraction. Moreover, sample sizes were typically smaller, and in some cases artifacts associated with concurrent DBS may have been included (Airaksinen et al., 2015; Park et al., 2009; Salenius et al., 2002).

After controlling for these shortcomings, our results revealed a possible relationship between the magnitude of CMC and the clinical symptoms in PD. The CMC proved to be a particularly sensitive measure. Interestingly, there was no relationship between clinical symptoms of PD and mean power in the beta range, despite a clear reduction of beta power in PD compared to control participants (Bosboom et al., 2006; Stoffers et al., 2007; Vardy et al., 2011). Furthermore, the relationship between CMC and motor impairment in participants remained even after controlling for changes in beta power. Thus, it is possible that our task and the CMC, by loading on relevant pathways (Baker, 2007; Tatsuya Mima et al., 2002), provide a more sensitive and objective measure of PD-related motor deficits, compared to beta power (see also (Proudfoot et al., 2018). The simple and well-controlled gripper task employed here, in conjunction with MEG (or EEG) and EMG recordings, may therefore provide a convenient and effective set-up in which to obtain a sensitive CMC marker to monitor disease progression or when testing the influence of novel treatments for PD. It will be interesting to investigate whether changes in cortico-muscular coupling are already present in early stages of the disease or in individuals at risk of developing PD (such as individuals with Rapid Eye-movement Sleep behavioral Disorder), thereby possibly providing a valuable early marker aiding risk assessment, stratification, diagnosis, and disease prognosis.

## Acknowledgements

We thank the volunteers and the NIHR National BioResource (https://bioresource.nihr.ac.uk/) which supported the recalling process of the volunteers. This work was funded by the Wellcome Trust (104571/Z/14/Z to KN and 098282/Z/12/Z to MH), and the British Academy (NZ), and supported by the National Institute for Health Research (NIHR) based at Oxford University Hospitals NHS Trust, and the NIHR Oxford Health Biomedical Research Centre. The Wellcome Centre for Integrative Neuroimaging is supported by core funding from the Wellcome Trust (203130/Z/16/Z).

## Supplementary Material

**Figure S1.**
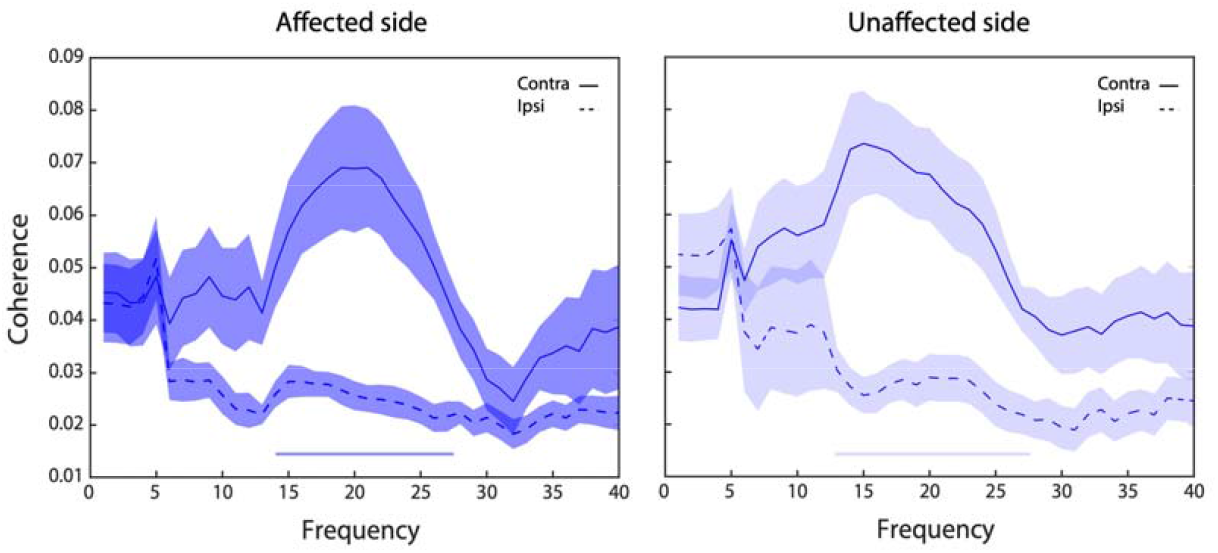
Cortico-muscular coherence (CMC) between the forearm EMG and contralateral (solid lines) and ipsilateral (dashed lines) motor cortex to the affected (darker blue – left graph) and unaffected (lighter blue – right graph) sides during the 1-3 s stable-grip period (shaded areas in Fig. 1b-c) in fifteen participants with Parkinson’s disease with tremor present in one hand as measured by questions 3.15, 2.16 and 3.17 in the UPDR III. Contralateral beta CMC was significantly higher than ipsilateral beta CMC for both affected and unaffected sides. There was no significant difference in contralateral CMC for the affected and unaffected sides.

**Figure S2.**
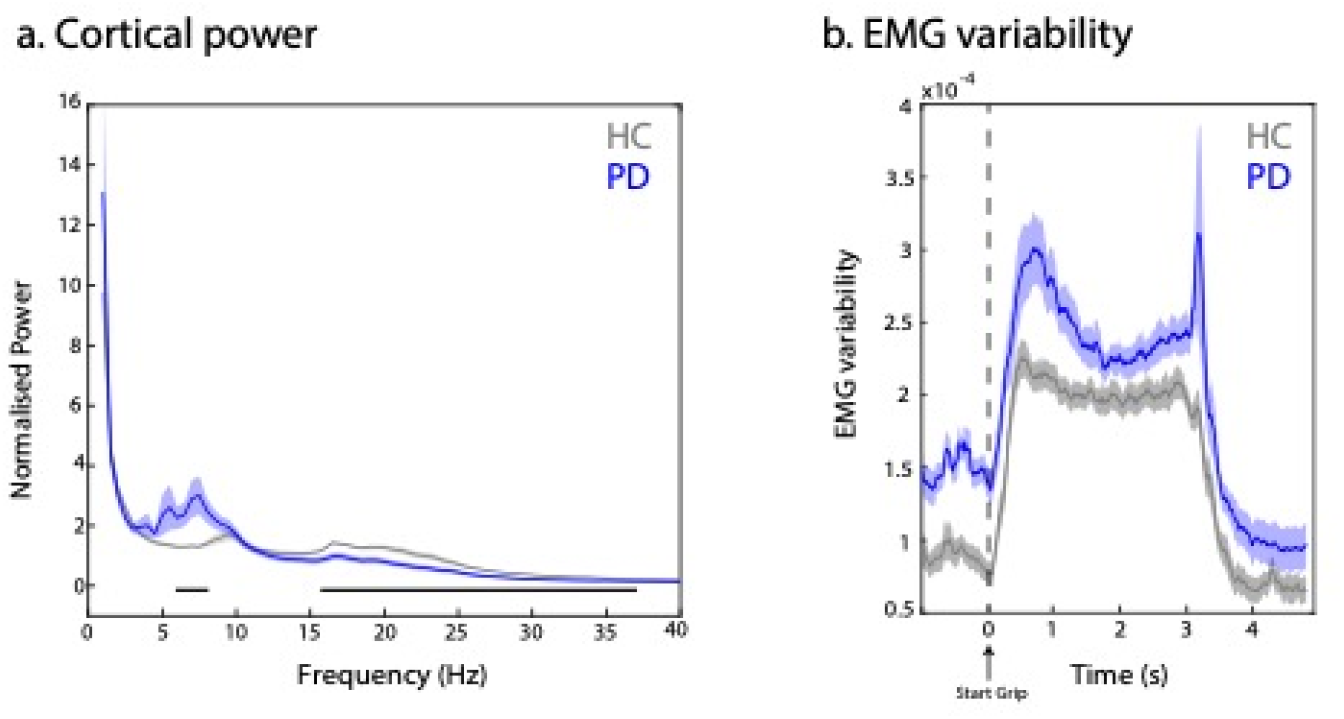
a) Normalised power (power at a given frequency divided by the average power at all other frequencies) in the pre-defined motor channels during the task. In line with previous studies, participants with PD (blue) had significantly stronger alpha power (6-8 Hz, cluster *p*= 0.05) and reduced beta power (15.5-37 Hz, cluster *p*= 0.0019). Mean beta power in PD participants was used as a covariate for any analysis examining the relationship between CMC and disease-related measures (see also **Figure S3**). It is important to note that even though changes to the alpha cluster in the PD group here may reflect tremor in these individuals (considering that only motor channels were used to obtain cortical power), similar changes in alpha power occur in non-motor channels in PD (Stoffers et al., 2007) and have also been reported in other neurogenerative conditions, such as Alzheimer’s disease (e.g. Osipova et al., 2004; Poza et al., 2004; Hughes et al., 2019). b) Participants with PD (blue) had significantly more variable EMG compared to healthy controls, though this was not restricted to the grip period. This variability may reflect tremor in some of these individuals. EMG variability at the time of the grip was used as a covariate for any analysis examining the relationship between CMC and disease-related measures (see also **Figure S3**).

**Figure S3.**
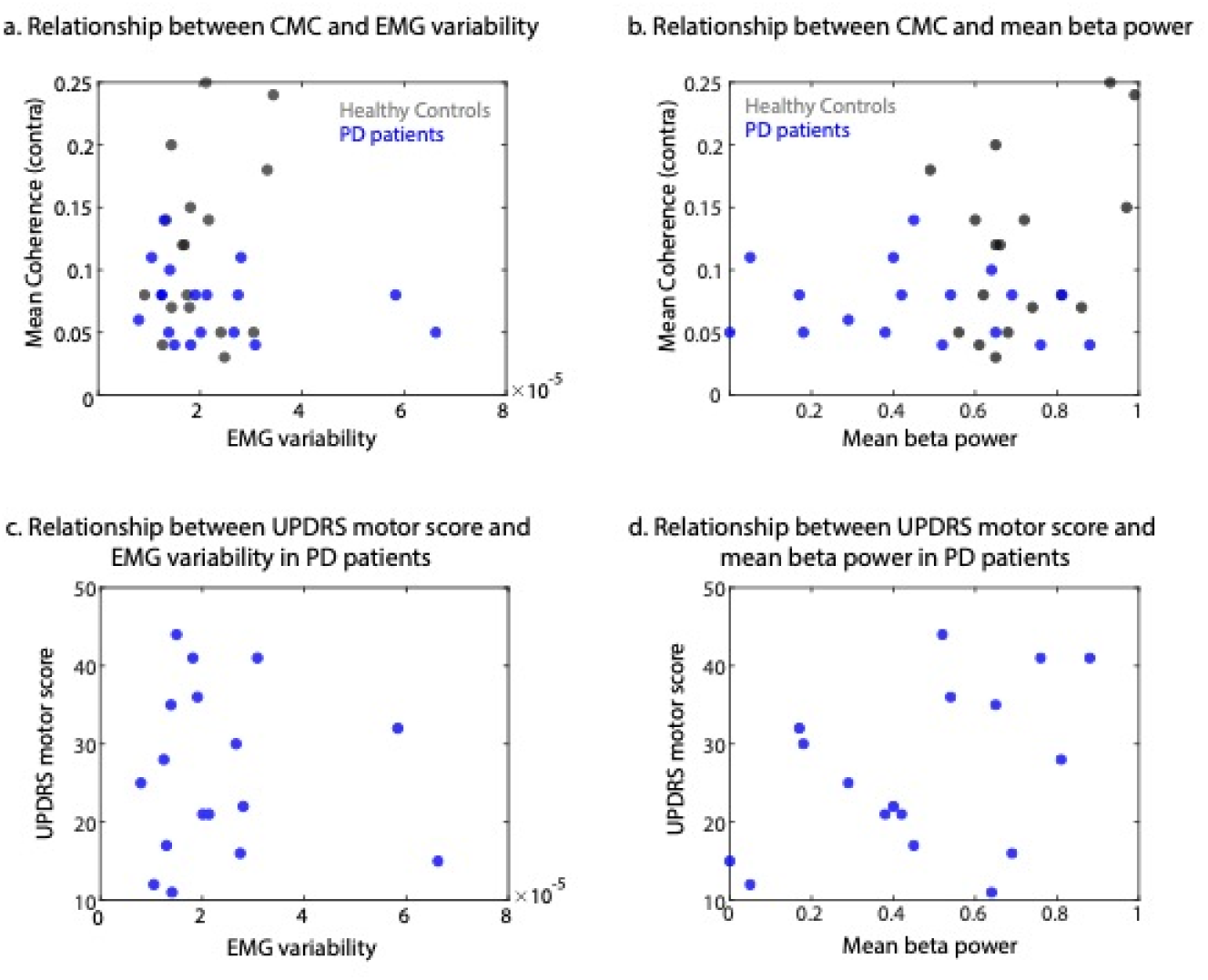
There were no relationships between CMC and EMG variability (both groups r= −0.010, *p*= 0.96, PD group only: r= −0.19, *p*= 0.46) (a), or between CMC and cortical beta power (r= 0.32, *p*=0.063) (b), across participants. In participants with PD, motor symptoms as measured by the UPDRS section III (motor symptoms), also did not correlate with EMG variability (r= −0.045, *p*= 863) (c) or with mean cortical beta power (r=0.43, *p*= 0.086) (d).

